# Transfusion of allogenic murine HOD red blood cells preferentially induces low-affinity, short-lived IgG antibodies that are germinal center independent

**DOI:** 10.1101/2025.01.16.633377

**Authors:** Jelena Medved, Abhinav Arneja, Tamara C. Moscovich, Neha Shah, Rachel J. Muppidi, Emily D. Burnett, Alexis R. Boscia, Benjamin N. Hester, Jayati Maram, Rishi Raghavan, Selby A. Ireland, Margaret B. Berberian, Elizabeth A. Stern-Green, Aditi S. Kodali, Conrad S. Niebuhr, Juan E. Salazar, Manjula Santhanakrishnan, Krystalyn E. Hudson, James C. Zimring, Jeanne E. Hendrickson, Chance John Luckey

## Abstract

Transfusion-induced anti-red blood cell (RBC) alloantibodies pose a significant risk to patients who require chronic transfusions. Anti-RBC alloantibodies can be remarkably short-lived (i.e. evanescent), leading to clinically relevant alloantibodies that are not detected in later pre-transfusion antibody screens. Subsequent transfusion of alloantigen-positive RBCs stimulates a rapid memory antibody response that may induce a delayed hemolytic transfusion reaction (DHTR), causing morbidity and occasional mortality in chronically transfused patients. It is unclear why transfusions favor evanescent antibody responses over long-lived antibodies typically observed upon infections and vaccinations. We therefore turned to the HOD mouse model of RBC alloimmunization to elucidate regulators of antibody persistence in response to allogenic transfusions. By following antibody responses over time in transfused mice, we found that HOD-specific alloantibodies rapidly decay within three months while vaccination-induced antibodies remain constant. Thus, the HOD model recapitulates RBC antibody evanescence. The rapid antibody evanescence suggests that transfusion is a poor inducer of germinal centers (GCs), specialized immunological structures where B cells differentiate into germinal center B (GC B) cells and undergo iterative rounds of affinity maturation, ultimately differentiating into long-lived plasma cells that can produce antibodies for decades. Consistent with this hypothesis, we failed to observe an increase in GC B cell formation in response to transfusion, and the majority of anti-RBC alloantibodies were low affinity when compared to vaccination. To formally test the functional requirement for GCs in anti-RBC alloantibody production, we employed two orthogonal approaches to disrupt GC formation: i) day 4 CD40L blockade and ii) genetic disruption of the GC-transcription factor BCL6 selectively in B cells. Both approaches fully blocked GC formation, yet anti-RBC alloantibody production was unchanged. Collectively, our data demonstrate that anti-HOD RBC alloantibodies are GC-independent, low affinity and short-lived. The GC-independence of HOD RBC IgG responses has important implications for understanding the cellular and molecular pathways that regulate the humoral immune response to transfused RBCs, potentially explaining anti-RBC alloantibody evanescence patterns in patients.

## Introduction

Red blood cell (RBC) transfusion from allogenic donors is a critical, life-saving treatment for patients with hematologic diseases, providing healthy functional RBCs to patients who have difficulty in producing their own. Despite the clinical utility of transfusion, the immune response to allogenic RBCs can cause significant problems for patients who require chronic transfusions^1,2^. Transfused RBCs typically express multiple antigens on their surface that are polymorphic between patients and donors. Exposure to these RBC alloantigens via transfusion can induce anti-RBC alloantibody production in recipients. Anti-RBC alloantibodies can make finding compatible RBCs difficult, limiting therapeutic options.

RBC alloantibodies generated in response to transfusion are different from antibodies formed in response to infection or vaccination. Transfusion-induced RBC alloantibodies are remarkably short-lived, as roughly a third of all anti-RBC antibodies become undetectable (evanescent) within a year of their generation in the general population^3,4^, while 60-80% of anti-RBC alloantibodies are evanescent in patients with sickle cell disease^5,6^. This stands in stark contrast to the antibody responses induced by vaccines and typical infections, where antigen-specific antibodies have half-lives that range from tens to hundreds of years^7^. The rapid evanescence of anti-RBC alloantibodies can be a major problem for patients; standard blood screening tests performed at the time of future transfusions may fail to detect previously generated alloantibodies. As a result, patients are at risk of receiving incompatible RBCs, re-exposure to which may lead to rapid induction of alloantigen-specific antibodies presumably through the activation of memory B cells and their differentiation into antibody secreting plasma cells. The rapid production of RBC-specific IgGs can destroy transfused RBCs and drive widespread systemic immune activation in a process referred to as a delayed hemolytic transfusion reaction (DHTR)^8,9^, a major cause of morbidity and mortality in chronically transfused patients.

DHTRs result from two key immunological phenomena that appear to be in direct contrast to our classical understanding of the basic immunological mechanisms that typically guide long-lived antibody production and B cell memory. In response to both vaccination and viral infections, long-lived plasma cells capable of secreting antibodies for decades as well as high-affinity memory B cells are generated in specialized immunological structures called germinal centers (GC) that are formed within the B cell follicle^10^. GCs support multiple rounds of T-B cell interactions leading to B cells undergoing somatic hypermutation and affinity maturation, selecting B cell clones which express high-affinity antibodies. GCs further induce the differentiation of these high-affinity clones into both long-lived memory B cells as well as long-lived plasma cells. Memory B cells do not actively secrete antibodies, but rather persist in secondary lymphoid organs awaiting re-exposure to the same antigen. Long-lived plasma cells on the other hand travel to the bone marrow where they can live for decades actively secreting antibodies. Many transfused and alloimmunized patients appear to have unlinked these two key GC functions in that anti-RBC specific antibody responses are short-lived yet memory/recall responses remain intact. We therefore set out to directly investigate the role of GC using a mouse model of transfusion-induced RBC-specific alloimmunization.

## Methods

### Mice

C57BL/6J WT, FVB WT, BCL6-floxed^11^ and CD19-Cre^12^ mice were purchased from Jackson Laboratories. BCL6-floxed mice were crossed to CD19-Cre to generate BCL6^fl/fl^CD19-Cre^+/-^ (BCL6-BKO) mice. HOD transgenic mice express a triple fusion construct of Hen Egg Lysozyme, Ovalbumin, and the human Duffy red blood cell antigen selectively on the surface of RBCs, and were generated on the FVB background as described previously^13^. All mice were bred and maintained at the animal facilities of the University of Virginia and Yale University. They were immunized between the ages of 8-10 weeks. Experimental groups were age and gender matched. All procedures were approved by Institutional Animal Care and Use Committees at the University of Virginia and at Yale University.

### Murine blood collection

Collection, processing, and storage of HOD blood was performed as described previously^14^. Briefly, blood from HOD mice was collected aseptically directly into the anticoagulant citrate phosphate dextrose adenine solution (CPDA-1, Boston Bioproducts IBB-420) by cardiac puncture. The final CPDA-1 concentration was kept at 20% (v/v). Collected HOD blood was leukoreduced using a whole blood cell leukoreduction filter (Pall, AP-4851). Leukoreduced blood was centrifuged at 1200 X *g* for 10 min, adjusted to a final hematocrit level of 75% through removal of supernatant, and stored at 4°C for 12 days before transfusion.

### HEL-OVA conjugation

HEL-OVA conjugation was done as previously described (Robson, 2008). Briefly, Ovalbumin (450 mg; Sigma A5503) and HEL (126 mg; Sigma L4919) were dissolved in 18 ml phosphate buffer (pH 7.5). The solution was centrifuged at 450 X g for 5 min and 14.4 ml of 0.3% glutaraldehyde solution in phosphate buffer (v/v) (Electron Microscopy Sciences 16120) was added to the supernatant and mixed for 1 h at room temperature before centrifugation at 450 X g for 5 min. The supernatant was then dialyzed (ThermoFisher 87737) overnight against PBS at 4°C before sterile filtration through 0.2 μm filters (ThermoFisher 564-0020). Protein content was determined using a Micro BCA Protein Assay Kit (ThermoFisher 23235) and aliquots of 1 mg/mL were stored at -20°C until required.

### Immunization and phlebotomy of mice

HOD blood was taken out of storage, pipetted up and down gently to make a uniform suspension of RBCs, and transfused into recipient mice via retro-orbital injection. Each recipient mouse received 100 μl of 12-day stored HOD blood, which is the volume adjusted equivalent of 1 unit of packed RBCs in humans. For the vaccination experiments, 100 μg of HEL-OVA conjugate was thoroughly mixed with 100 μl aluminum hydroxide (ALHYDROGEL, vac-alu-250, InvivoGen) and given to recipient mice via intraperitoneal (IP) injections. Blood samples for antibody measurements from immunized mice were collected by submandibular vein puncture and allowed to clot. Samples were then spun at 5,000 rpm for 5 min and the supernatant was transferred to new tubes. This process was then repeated for the second time in order to get sera with no RBC contamination. Sera were stored at -20°C until analysis for anti-RBC alloantibodies.

### Antibody treatment

For CD40L blocking experiments, mice were given 250 μg of the anti-mouse CD40L blocking antibody or isotype control via IP injection (BioXCell, BE0017-1 or BE0091) on days 4, 7, 10, and 14 post-immunization.

### Antibody detection by ELISA

To eliminate day-to-day technical variability, all sera samples from a single experiment were batched, run and analyzed on the same day. Anti-HEL responses were measured by HEL-specific enzyme-linked immunosorbent assay (ELISA) as previously described^14,15^. Briefly, high binding polystyrene plates (Corning 9018) were coated overnight at 4ºC with 10 μg/ml HEL (Sigma-Aldrich) in PBS. Plates were then washed with wash buffer (0.05% Tween-20 in PBS) and incubated with blocking buffer (2% BSA and 0.05% Tween-20 in PBS). Serum samples were serially diluted (starting at 1:25) in blocking buffer and incubated in coated plates for 1 hour at room temperature. Plates were then washed 3 times and horseradish peroxidase-conjugated goat anti-mouse IgG Fcγ specific antibody (Jackson ImmunoResearch) was then used as a secondary stain at a 1:5,000 dilution. After washing the plates twice with wash buffer and 4 times with PBS, wells were developed using TMB substrate (SeraCare) and quenched with 2N H_2_SO_4_ after 10 min at room temperature. Optical densities were measured at 450 nm. End-point titers were calculated using GraphPad Prism through interpolation of the cutoff value from the fit of the optical density vs. (1/serum dilution) curve for each sample using the “plateau followed by one-phase exponential decay” model. The cutoff value was defined as the average plus three standard deviations of signals from background wells (i.e. signal values from wells incubated with blocking buffer alone). For ELISA assays that employed a urea wash, 6.5M urea in DI water was added for 10 minutes after the incubation with sera samples and before the secondary antibody was added. Plates were washed twice before and after incubation with urea.

### Germinal center B cell detection by flow cytometry

Spleens were harvested and dissociated using gentle mechanical disruption. Single cell suspension was subjected to RBC lysis with RBC lysis buffer (ThermoFisher), resuspended in growth medium (10% FBS in RPMI) and filtered through 40 μm cell strainers (Falcon). 5 million cells were used for staining. All staining antibodies were diluted in FACS buffer (2% FBS, 0.5% BSA, and 0.5% NaN_3_ in PBS). GC B cells were characterized and enumerated by flow cytometry. GC B surface staining was performed for: Brilliant Violet 421™ anti-mouse CD19 (BioLegend), Brilliant Violet 711™ anti-mouse IgD (BioLegend), Fluorescein Peanut Agglutinin (Vector Laboratories) and PE anti-mouse/human GL7 antigen (BioLegend). Brilliant Violet 605™ anti-mouse CD4 (1:400), Brilliant Violet 605™ anti-mouse CD8 (1:400) and dead cells (Zombie Yellow Dye™, BioLegend) were excluded in a dump channel. To determine the expression of BCL6, splenocyte surface staining was first performed as described above, followed by fixing and permeabilization using the Transcription Factor Buffer Set (BD Pharmingen). Intracellular staining was performed with either Alexa Fluor 647 conjugated mouse anti-BCL6 antibody (BD Pharmingen) or Alexa Fluor 647 mouse IgG1κ isotype control (BD Pharmingen) was used to determine the unspecific staining. Flow cytometry was performed using the Attune Nxt and data were analyzed using FlowJo and GraphPad Prism.

### Statistical analysis

Statistical analyses and graphing were performed with GraphPad Prism software. Groups of interest were compared using Mann-Whitney U tests, preceded with Kruskal-Wallis tests where applicable. Antibody evanescence time-course data were compared using the Friedman test for repeated measures, followed by post-hoc Dunn’s test to compare individual groups of interest. A value of P < 0.05 was considered to be statistically significant and assigned *, whereas P < 0.01, P < 0.001 and P < 0.0001 were assigned **, *** and ****, respectively.

## Results

### Transfusion-induced IgG levels decay more rapidly than those induced by vaccination

To determine if RBC transfusion preferentially induced evanescent alloantibodies, we compared the longevity of antibody responses generated to RBC transfusion vs. protein-adjuvant vaccination. A well-established mouse model for the study of transfusion mediated RBC alloimmunization, HOD mice express exclusively on RBCs a transgenic chimeric protein that consists of the human Duffy antigen fused to a portion of chicken ovalbumin and the entire hen egg lysozyme (HEL) protein^13^. We compared antibody responses to HOD RBC transfusion with antibody responses induced by conjugated HEL-OVA proteins emulsified in Aluminum salts (HEL-OVA/Alum) (Figure 1A). Transfusion of HOD RBCs induced an anti-HEL specific IgM response that peaked at 1 week (data not shown), while the anti-HEL IgG response peaked at 2 weeks. Anti-HEL IgG titers in transfused mice rapidly declined over the course of 13 weeks (Figure 1B). In contrast, mice immunized with HEL-OVA/Alum generated anti-HEL IgG that remained stable over the same period (Figure 1C). Thus, the HOD RBC transfusion model recapitulates rapid evanescence observed in some patients.

**Figure 1.**
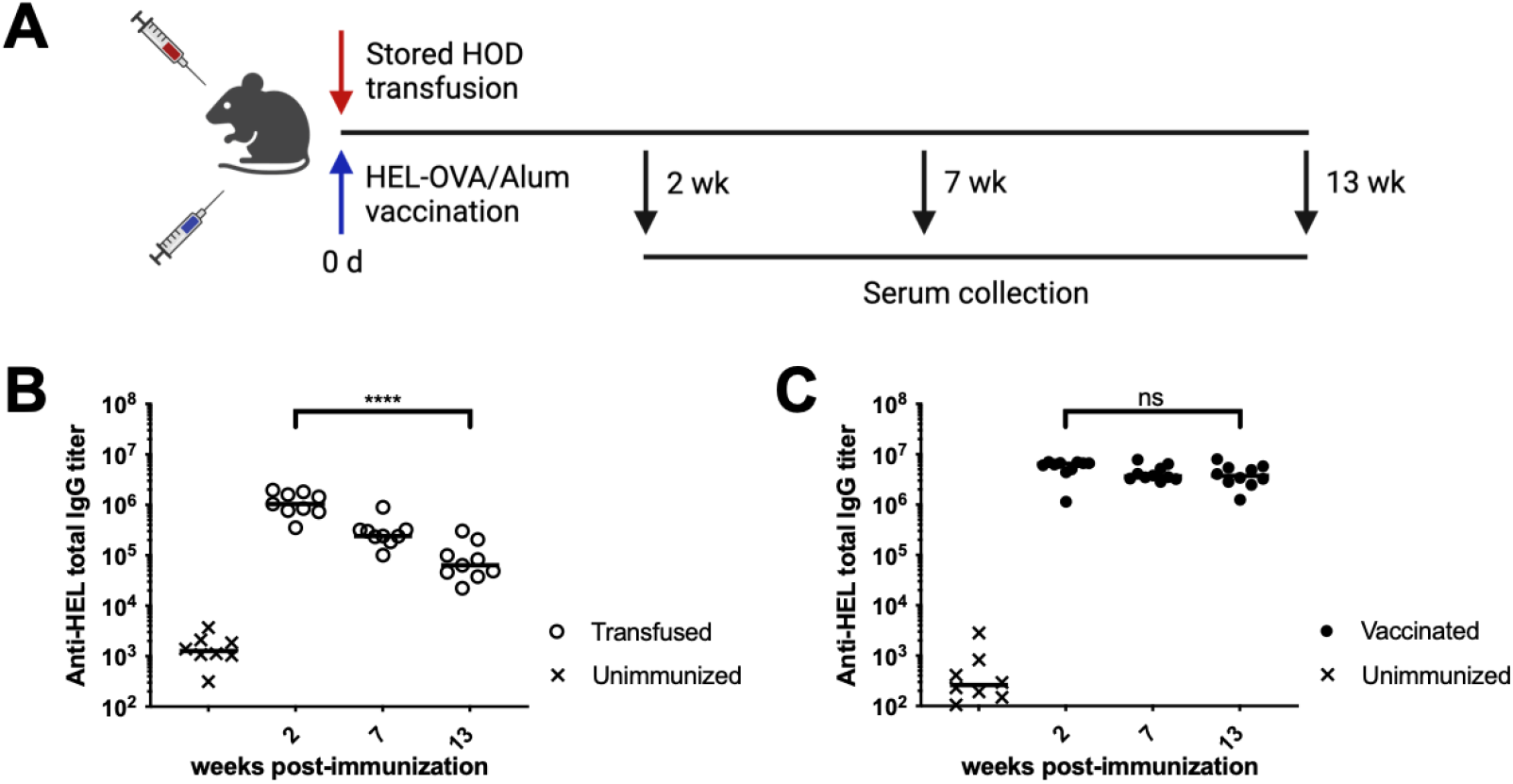
Transfusion-induced IgG levels decay more rapidly than those induced by vaccination. (A) Experimental approach. WT mice were either transfused with HOD RBCs or vaccinated with HEL-OVA/Alum. On weeks 2, 7 and 13, sera were collected for assessment of IgG levels by limiting dilution ELISA. (B) IgG titers of transfused mice. (C) IgG titers of vaccinated mice. Each data point represents one mouse. Bars on scatter plots are median values. Figure shows are representative of 4 independent experiments. Groups of interest were compared using the non-parametric Friedman test for repeated measures, followed by post-hoc comparisons of individual groups using Dunn’s test. *P<0.05, **P<0.01, ***P<0.001, ****P<0.0001, ns P>0.5.

### Transfusion fails to induce robust Germinal Center B cell formation

Given the established role of GCs in the generation of long-lived antibodies^10^, we investigated whether transfusion of HOD RBCs leads to less efficient differentiation of GCs compared to HEL-OVA/Alum vaccination. As the spleen plays a central role in alloantibody production in response to HOD RBC transfusion^16^, we concentrated analysis on splenic immune cell populations. To unambiguously identify and quantify GC responses, GC-localized B cells were identified and quantified by flow cytometry based on surface staining pattern of B cell markers (CD19^+^IgD^lo^PNA^hi^GL7^hi^) plus intracellular expression of the GC-specific transcription factor BCL6^10^ (gating strategy is shown in Supplemental Figure 1). Spleens were collected 2 weeks post-immunization, corresponding to both peak IgG response and peak GC reaction in response to vaccination (Figure 2A)^7^. As expected, vaccinated mice showed strong induction of splenic GC B cells, reflecting a robust GC response (Figures 2B and 2C). In contrast, transfused mice exhibited no increase in splenic GC B cells above levels from naive mice (Figures 2B and 2C). Thus, unlike HEL-OVA/Alum vaccination, transfusion with HOD RBCs does not trigger a robust GC response.

**Figure 2.**
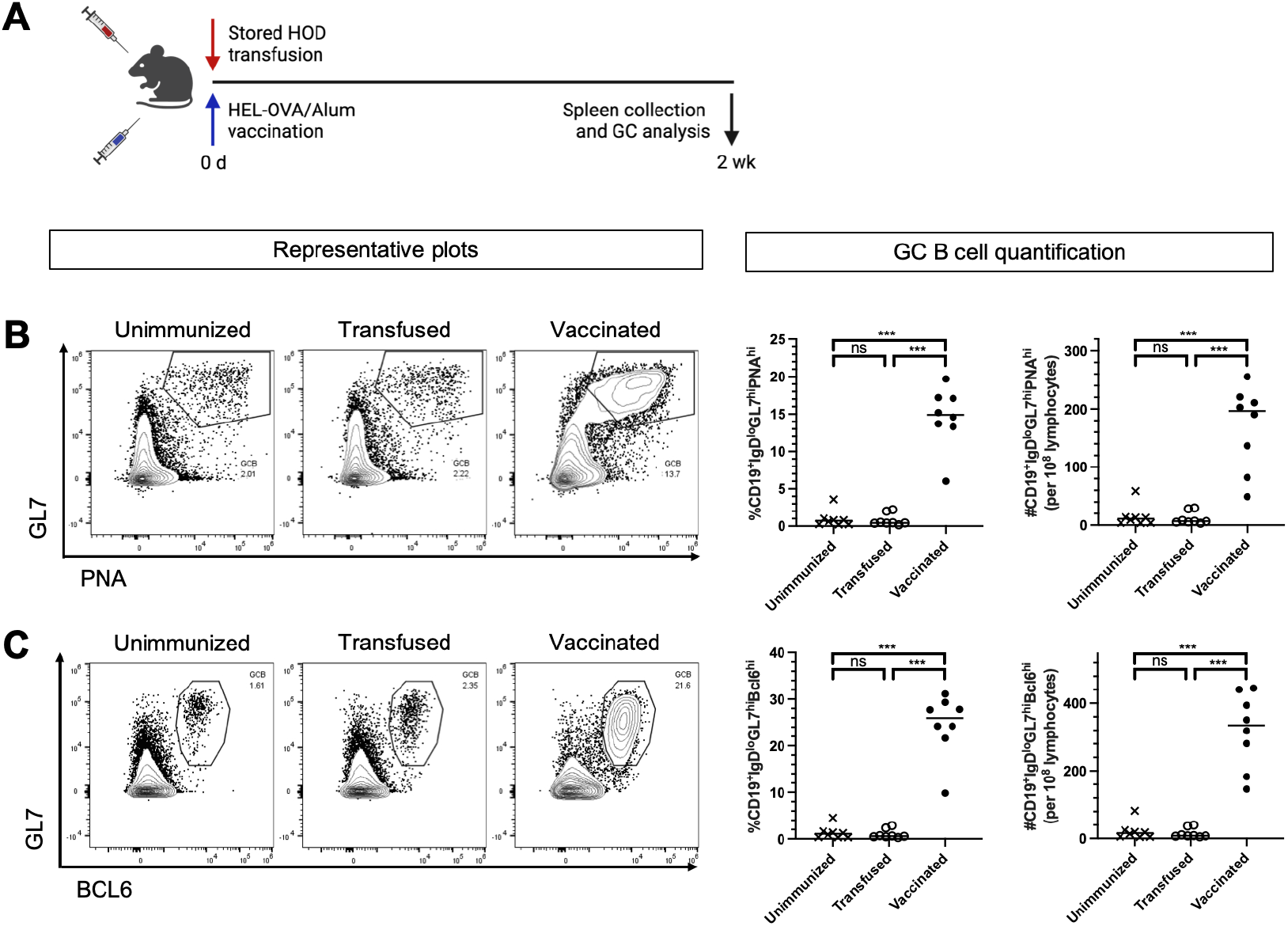
Transfusion fails to drive robust GC formation. (A) Experimental approach. WT mice were either transfused with HOD RBCs or vaccinated with HEL-OVA/Alum. Spleens were collected 2 weeks after immunization and the production of splenic GC B cells was analyzed by flow cytometry. Representative flow plots and GC B cell quantification for surface (B) and intracellular (C) staining. Each data point on scatter plots represents one mouse. Bars on scatter plots are median values. Data are representative of 4 independent experiments. Groups of interest were compared using Mann-Whitney U tests, preceded with Kruskal-Wallis tests. *P<0.05, **P<0.01, ***P<0.001, ****P<0.0001, ns P>0.5.

### Anti-RBC IgG antibodies are mostly low affinity

In addition to their role in generation of long-lived antibody responses, GCs are critical for B cell affinity maturation. Affinity maturation is a process by which antigen-specific B cells undergo multiple rounds of mutation and selection, leading to higher affinity antibodies. To directly measure antigen-specific antibody affinity, we used a modified ELISA protocol where antigen-bound primary antibodies were washed with 6.5M urea prior to addition of secondary antibody, an experimental maneuver that selectively removes low-affinity antibodies but not high affinity antibodies^17^. We collected sera from recipient mice 2 weeks post-immunization with either transfusion or vaccination and compared antibody levels with or without urea wash (Figure 3A). Without urea treatment, ELISA revealed robust IgG titers in sera from transfused mice, (Figure 3B). Treatment with urea led to a 100-fold reduction in IgG titers, demonstrating that the majority of transfusion generated IgGs are low affinity. In contrast, vaccinated sera showed a more modest reduction in IgG titers (less than 5-fold) following urea treatment (Figure 3C), reflecting higher affinity IgGs.

**Figure 3.**
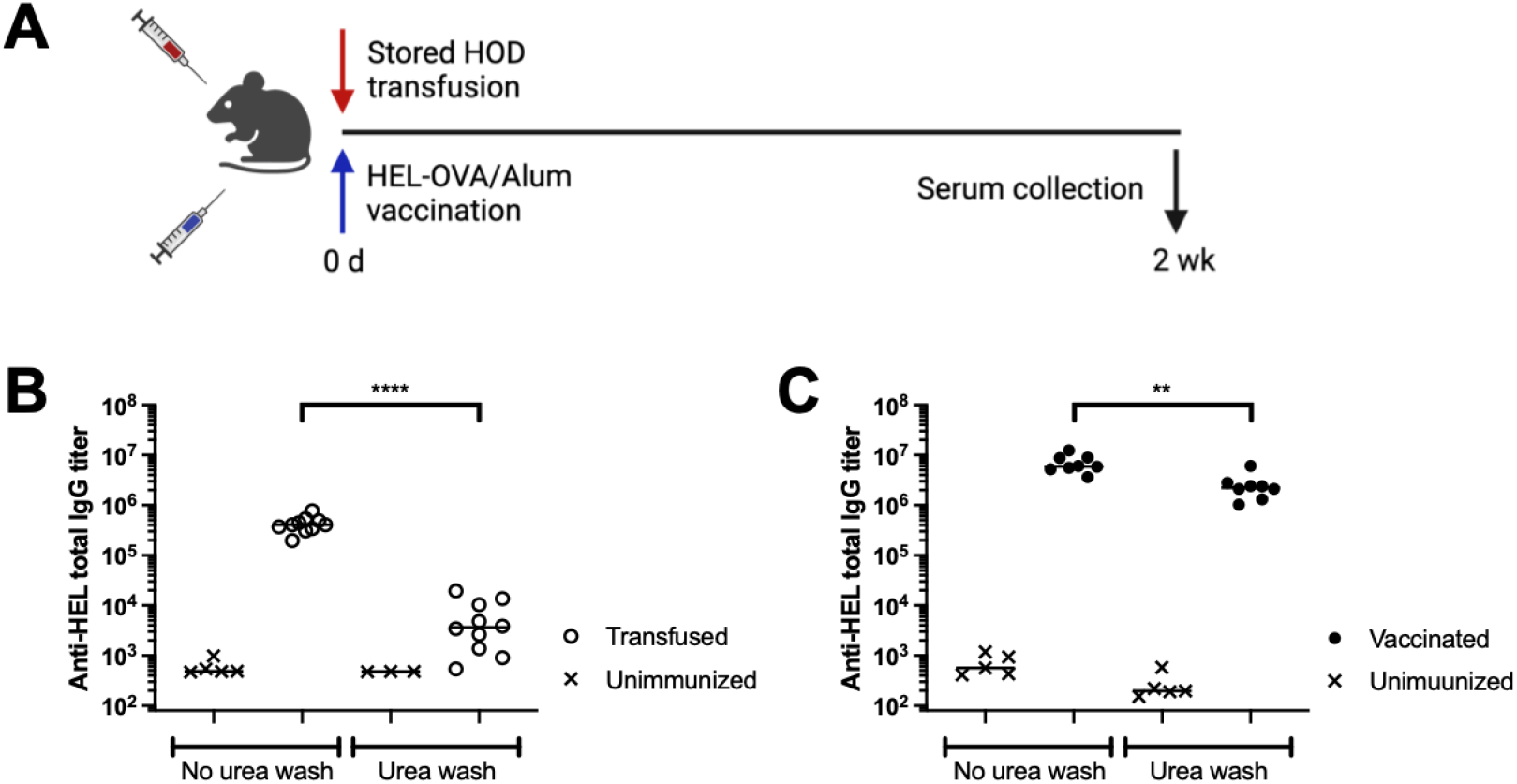
Most anti-RBC IgG antibodies are low affinity. (A) Experimental approach. WT mice were either transfused with HOD RBCs or vaccinated with HEL-OVA/Alum. Sera were collected 2 weeks after immunization for assessment of IgG levels by limiting dilution ELISA that either included or did not include urea wash. (B) IgG titers of transfused mice. (C) IgG titers of vaccinated mice. Each data point represents one mouse. Bars on scatter plots are median values. Data are representative of 4 independent experiments. Groups of interest were compared using Kruskal-Wallis test followed by post-hoc Mann-Whitney U tests. *P<0.05, **P<0.01, ***P<0.001, ****P<0.0001, ns P>0.5.

### Disruption of GC formation does not alter anti-RBC IgG antibody production

To formally investigate the functional role of GCs in anti-RBC alloantibody responses, we used mice with disrupted GCs. GC responses rely on prolonged interactions between antigen-specific T cells and antigen-specific B cells, which in turn depend on CD40L from helper CD4^+^ T cells engaging with CD40 on B cells^10^. Administration of CD40L blocking antibody to recipient mice starting 4 days after the immune stimulus allows for initial T-B cell interactions and class-switching, but selectively blocks the long-lived T-B interactions required for GC formation^18^. We therefore treated transfused mice or vaccinated mice with CD40L blocking antibody on days 4-10 post-immunization (Figure 4A). Flow cytometry demonstrated near complete ablation of all GC B cells in CD40L-blocked mice (Supplemental Figure 2). Day 4 CD40L blockade led to a significant reduction in HEL-specific IgG levels in vaccinated mice (Figure 4B), but not in transfused mice (Figure 4C), supporting the hypothesis that GCs play only a minor role in anti-RBC alloantibody responses to HOD RBC transfusion.

**Figure 4.**
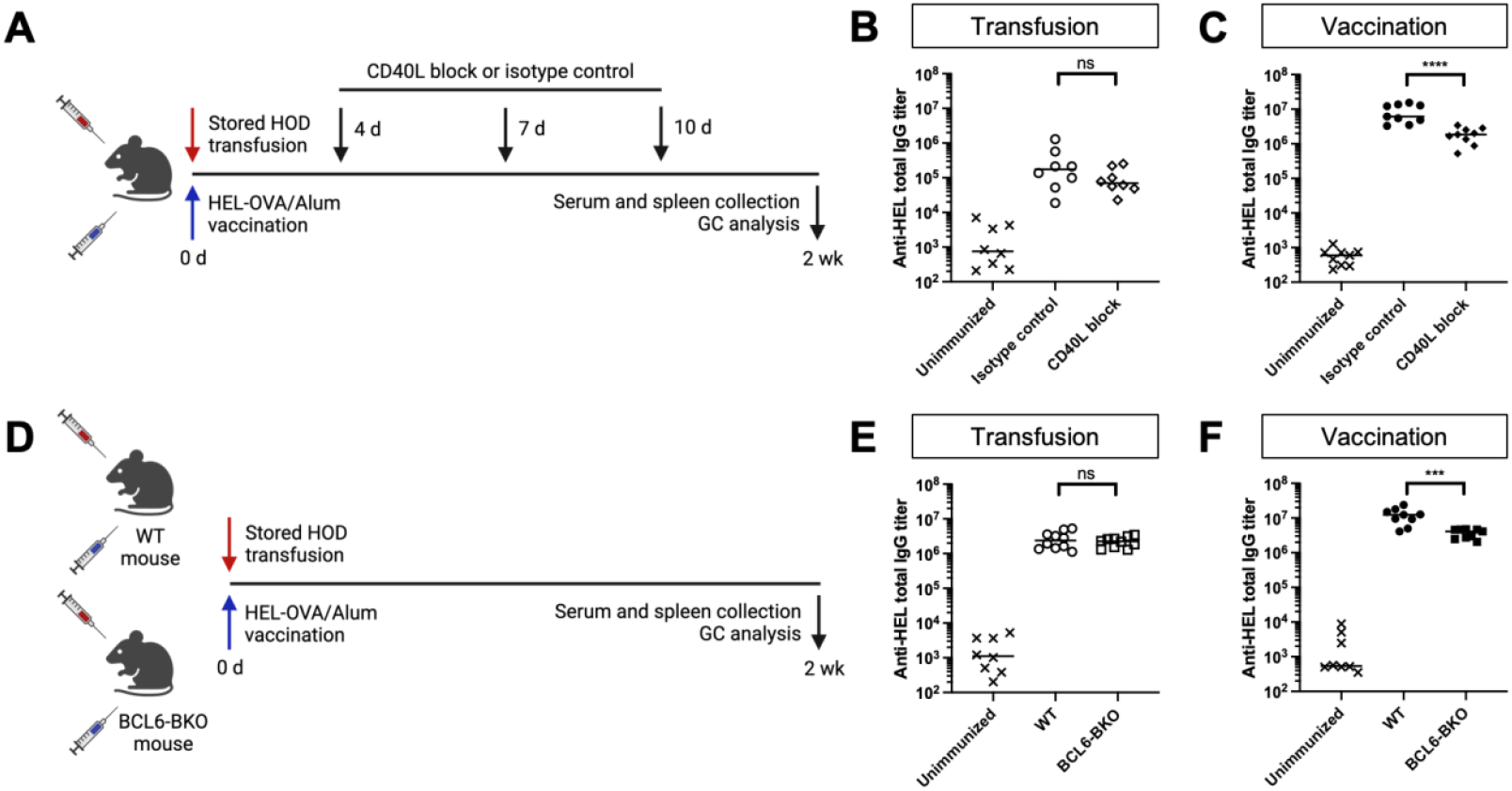
Disruption of GC formation does not alter anti-RBC IgG antibody production. (A) Experimental approach for CD40L blocking experiments. WT mice were either transfused with HOD RBCs or vaccinated with HEL-OVA/Alum. They received CD40L blocking antibody 4 days after immunization and subsequently on days 7 and 10 pi. Sera were collected 2 weeks after immunization for assessment of IgG levels by limiting dilution ELISA. (B) IgG titers of transfused mice. (C) IgG titers of vaccinated mice. (D) Experimental approach for experiments that utilized BCL6-BKO mice. WT or BCL6-BKO mice were either transfused with HOD RBCs or vaccinated with HEL-OVA/Alum. Sera were collected 2 weeks after immunization for assessment of IgG levels by limiting dilution ELISA. (E) IgG titers of transfused mice. (F) IgG titers of vaccinated mice. Each data point represents on mouse. Bars on scatter plots are median values. Figure shows a representative experiment out of 3. Groups of interest were compared using Mann-Whitney U tests preceded by Kruskal-Wallis tests. *P<0.05, **P<0.01, ***P<0.001, ****P<0.0001, ns P>0.5.

We independently tested the functional role of GCs using an orthogonal approach to disrupt GC formation. Endogenous expression of the transcription factor BCL6 in B cells is essential for the development of GCs^19^. Mice with conditional B-cell-specific knockouts of BCL6 were generated (BCL6-BKO mice; see Methods) and had highly impaired GC development in response to either vaccination or transfusion (Supplemental Figure 2). BCL6-BKO were transfused or vaccinated as before (Figure 4D), and demonstrated responses similar to mice treated with CD40L blockade. Vaccinated BCL6-BKO mice expressed significantly lower antigen-specific IgG compared to control mice, whereas transfused BCL6-BKO mice had levels of antigen-specific IgG similar to controls (Figures 4E and 4F). Thus, preventing GC formation either by day 4 CD40L blockade or by BCL6 deletion in B cells does not significantly impact IgG production following transfusion. We therefore conclude that most anti-RBC alloantibodies in response to transfusion are produced in a GC-independent manner.

## Discussion

Anti-RBC alloantibodies generated in response to HOD RBCs are different from those generated in response to the cognate antigen emulsified in an Alum adjuvant. Specifically, transfusion of RBCs expressing the allogenic HOD antigen are poor inducers of GCs, resulting in the preferential production of low-affinity, short-lived IgG antibodies. Thus, the HOD transfusion model mimics antibody evanescence observed in many patients. Rapidly evanescent patient anti-RBC alloantibodies are unlikely to have undergone affinity maturation, and may represent predominantly low-affinity antibodies. While current clinical approaches are not optimized for measuring anti-RBC affinity, washing of RBCs incubated with patient sera with chaotropes prior to measuring agglutination reactions could allow for the measurement of affinity changes in patient responses over time.

Though most transfusion-induced anti-HOD alloantibodies are produced outside of GCs, it remains unclear where anti-HOD antibody secreting plasmablasts are located. Anti-RBC alloantibodies induced by HOD transfusion are produced predominantly by B cells initially localized in the marginal zone rather than by follicle B cells^20^. Classical marginal zone B cells (MZB) travel to the B cell follicle in a S1PR1 dependent manner, where they transfer their antigen to follicular B cells^21^. However, anti-HOD antibodies are unaffected in S1PR1^TSS^-KO mice, suggesting that MZB do not have to travel to the B cell follicle to generate anti-HOD antibodies. Since class-switching to IgG in the HOD system is clearly dependent on CD4^+^ T cells^20,22,23^, MZB must be interacting with HOD-specific T cells. However, it is unclear where interactions that drive IgG class-switching and plasmablast differentiation occur within the spleen. The mechanisms that govern CD4^+^ T-cell dependent MZB responses are poorly understood. The HOD model provides a unique opportunity to better understand how and where MZBs interact with helper CD4^+^ T cells to subsequently undergo class-switching and differentiation into IgG-secreting plasmablasts.

While anti-HOD antibodies are generated outside of GCs, it is unclear whether GC non-involvement is due to a lack of sufficient GC-inducing signals, or instead to an active suppression of GC formation in response to transfusion. We previously published that HOD transfusion induces a subset of HOD-specific CD4^+^ T cells to express both surface markers CXCR5 and PD1 and a transcription factor BCL6 consistent with their differentiation into “Pre-TFH” cells capable of providing initial B cell help for class-switching and plasmablast differentiation^15^. The lack of robust GC B cells in response to transfusion suggests that these transfusion-induced early “Pre-TFH” may be incapable of full differentiation into GC-localized “GC-TFH” ^24,25^. Though GC responses are readily induced by most vaccinations and infectious diseases, certain infections can actively inhibit GC production^26^. Prominent among these are antibody responses generated to malarial proteins specifically expressed during the blood stages of infection^27^. Malaria-infected RBCs actively suppress the production of GC through multiple mechanisms^26^, perhaps explaining why many children fail to generate high-affinity, long-lived protective antibodies to initial malarial infections, and remain susceptible to multiple rounds of re-infection.

Finally, our findings have important implications for understanding how long-lived memory B cells are generated in response to transfusion. Delayed hemolytic transfusion reactions occur because, despite evanescence, re-exposure through subsequent transfusion induces robust memory responses ^8,9^. This begs the question of where anti-RBC memory B cells are generated. Though the majority of memory B cells are thought to be generated in GCs, there is growing appreciation that B cells localized outside of GCs are also capable of differentiating into memory B cells^18,26^. That transfusion is a poor inducer of GCs suggests that anti-RBC memory B cells are generated from early MZB cells that have failed to undergo somatic hypermutation and affinity maturation. However, we cannot rule out the possibility that some GCs are activated at low levels in response to transfusion, and that anti-RBC memory cells arise from low-level GC responses. Future studies measuring recall responses in mice lacking GCs should help clarify whether GCs are required for anti-RBC memory B cell production.

While many anti-RBC alloantibodies are evanescent in patients, many other antibody responses appear stable over time. This suggests that there are multiple unknown factors that regulate whether a given transfusion can robustly induce GCs or not. One of the major limitations of this study is that it is limited to the HOD mouse model of RBC alloimmunization, and specifically to IgG responses induced by extended refrigerator storage of HOD RBCs. Further study of antibody persistence and affinity maturation in other antigen systems and in the context of other pro-inflammatory stimuli should provide key mechanistic information on the regulators of transfusion-induced GC responses.

## Notes

### Competing Interest Statement

The authors have declared no competing interest.

### Summary of Updates

The original manuscript was lacking funding sources which have now been added. No further edits were made.

### Funding

- NIH Common Fund: R01HL134691, P01HL169552

