## Supplemental Information for "Transfusion of allogenic murine HOD red blood cells preferentially induces low-affinity, short-lived IgG antibodies that are germinal center independent"

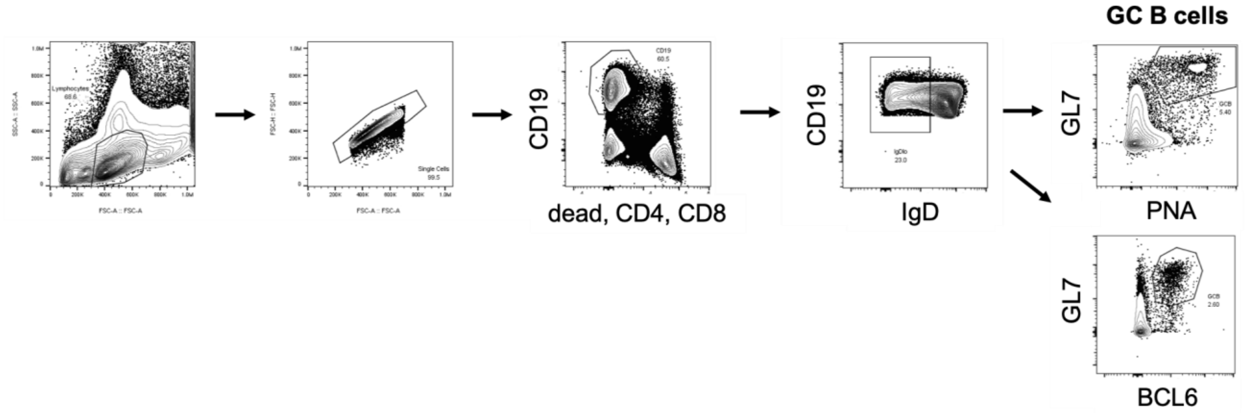


**Supplemental Figure 1. Gating strategy for identification of GC B cells.** Lymphocytes were first gated, followed by the exclusion of doublets. From the single cell population, CD19$^+$ cells were selected, while dead, CD4$^+$ and CD8$^+$ cells were excluded from the analysis. Activated B cells were distinguished based on the downregulation of IgD. GC B cells were characterized by two distinct strategies. For surface staining, GC B cells were identified based on the co-expression of high levels of GL7 and PNA. In parallel, an intracellular staining method was employed, where GC B cells were defined by the co-expression of GL7 and BCL6, a critical regulator of GC B cell differentiation and function.

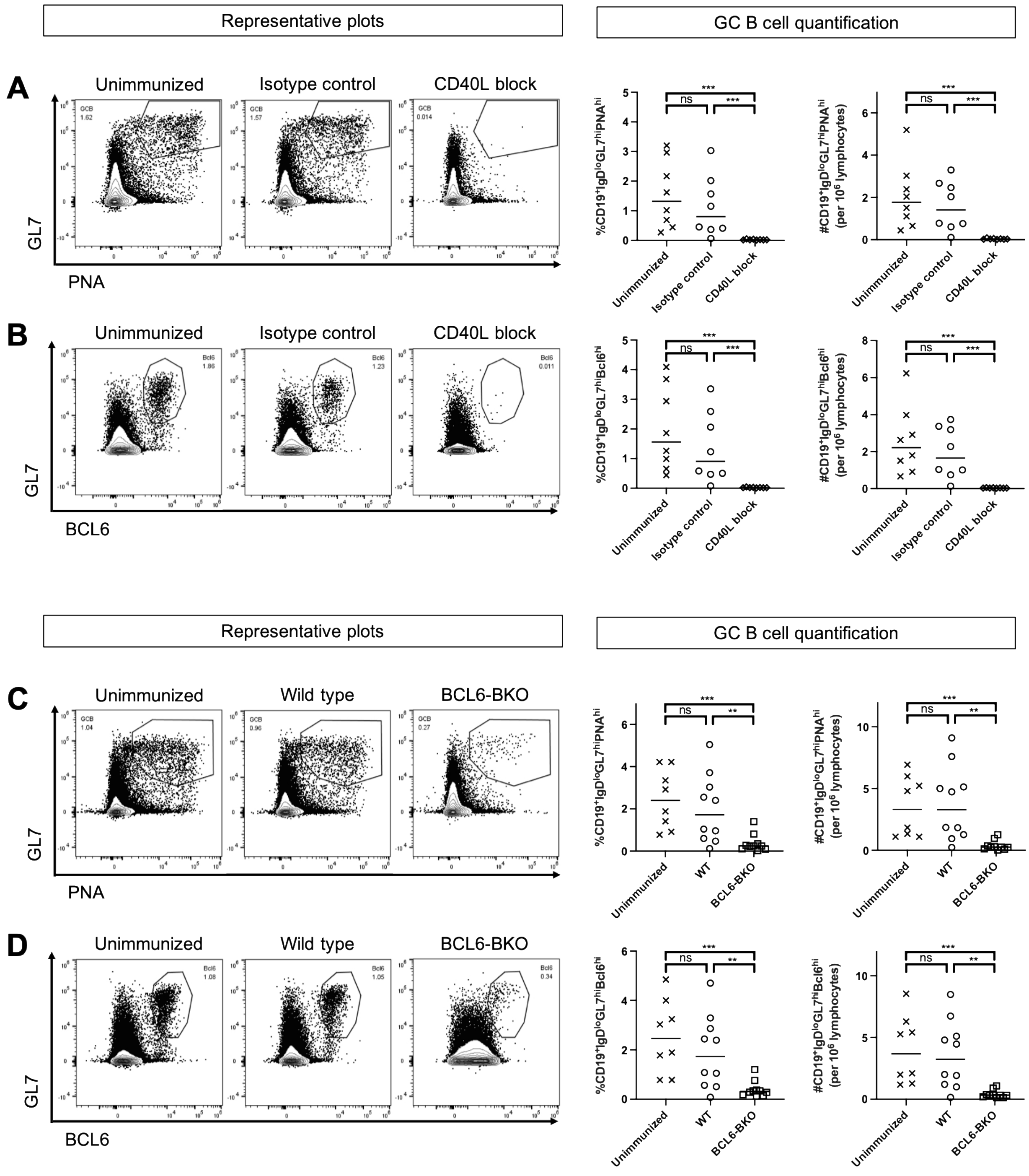


**Supplemental Figure 2. Confirmation of GC disruption.** Top. Representative flow plots of the surface (A) and intracellular (B) stain and GC B cell quantification in the mice that had GCs disrupted by injection of CD40L blocking antibody. WT mice were transfused and treated with either CD40L blocking antibody (CD40L block) or isotype control (Isotype control). Control WT (Unimmunized) mice were neither transfused nor treated with antibodies. Bottom. Representative flow plots of the surface (C) and intracellular (D) stain and GC B cell quantification in the mice that had GCs disrupted by disruption of BCL6 expression in B cells. WT or BCL6-BKO mice were transfused, while control (Unimmunized) mice were wild type mice that were not transfused. GC

B cell quantification is presented as GC B percent and number. Each data point on scatter plots represents one mouse. Bars on scatter plots are median values. Data are representative of 3 independent experiments. Groups of interest were compared using Mann-Whitney U tests, preceded with Kruskal-Wallis tests. *P<0.05, **P<0.01, ***P<0.001, ****P<0.0001, ns P>0.5.
